# A beneficial genomic rearrangement creates multiple versions of calcipressin in *C. elegans*

**DOI:** 10.1101/578088

**Authors:** Yuehui Zhao, Jason Wan, Shweta Biliya, Shannon C. Brady, Daehan Lee, Erik C. Andersen, Fredrik O. Vannberg, Hang Lu, Patrick T. McGrath

**Affiliations:** School of Biological Sciences, Georgia Institute of Technology, Atlanta, GA 30332, USA; The Wallace H. Coulter Department of Biomedical Engineering, Georgia Institute of Technology and Emory University, Atlanta, GA 30332, USA; Department of Molecular Biosciences, Northwestern University, Evanston, IL 60201, USA; Parker H. Petit Institute of Bioengineering and Bioscience, Georgia Institute of Technology, Atlanta, GA 30332, USA; School of Chemical & Biomolecular Engineering, Georgia Institute of Technology, Atlanta, GA 30332, USA; School of Physics, Georgia Institute of Technology, Atlanta, GA 30332, USA

## Abstract

Gene duplication is a major source of genetic novelty and evolutionary adaptation, providing a molecular substrate that can generate biological complexity and diversity (Ohno 1967, Taylor and Raes 2004). Despite an abundance of genomic evidence from extant organisms suggesting the importance of gene duplication, consensus about how they arise and functionally diversify is lacking (Innan and Kondrashov 2010). In the process of studying the adaptation of laboratory strains of C. elegans to new food sources, we identified a recombinant inbred line (RIL) with higher relative fitness and hyperactive exploration behavior compared to either parental strain. Using bulk-segregant analysis and short-read resequencing, we identified a de novo beneficial, complex rearrangement of the rcan-1 gene, which we resolved into five new unique tandem inversion/duplications using Oxford Nanopore long-read sequencing. rcan-1 encodes an ortholog to human RCAN1/DSCR1, which has been implicated as a causal gene for Down syndrome (Fuentes, Genesca et al. 2000). The genomic rearrangement in rcan-1 causes two complete and two truncated versions of the rcan-1 coding region, with a variety of modified promoter and 3’ regions, which ultimately reduce whole-body gene expression. This rearrangement does not phenocopy a loss-of-function allele, which indicates that the rearrangement was necessary for the observed fitness gains. Our results demonstrate that adaptation can occur through unexpectedly complex genetic changes that can simultaneously duplicate and diversify a gene, providing the molecular substrate for future evolutionary change.

---

Evolution is relentless, even in strains of model organisms used to study fundamental biological processes in laboratory settings. Despite experimenters creating environments filled with food and free of predators, animals adapt to their new and unnatural conditions (Yvert, Brem *et al*. 2003, Kasahara, Abe *et al.* 2010, Marks, Castro-Rojas *et al.* 2010, Duveau and Felix 2012, Orozco-terWengel, Kapun *et al.* 2012, Goto, Tanave *et al.* 2013, Large, Xu *et al.* 2016, Stanley and Kulathinal 2016, Zhao, Long *et al.* 2018). One example of laboratory adaptation is the standard strain of *C. elegans*, N2, which was isolated in 1951 from mushroom compost (**Figure 1a**). This strain spent almost two decades growing in the laboratory before methods of cryopreservation were developed (Sterken, Snoek *et al.* 2015). Mutations that arose and fixed in N2 after 1958 can be identified using a second lineage, called LSJ2, that separated from the N2 population at that time (McGrath, Xu *et al.* 2011). Approximately 300 mutations distinguish the N2 and LSJ2 strains. Previous studies have identified beneficial mutations that arose and fixed in the N2 lineage in the *glb-5*, *nath-10*, and *npr-1* genes (Duveau and Felix 2012, Zhao, Long *et al.* 2018).

**Figure 1.**
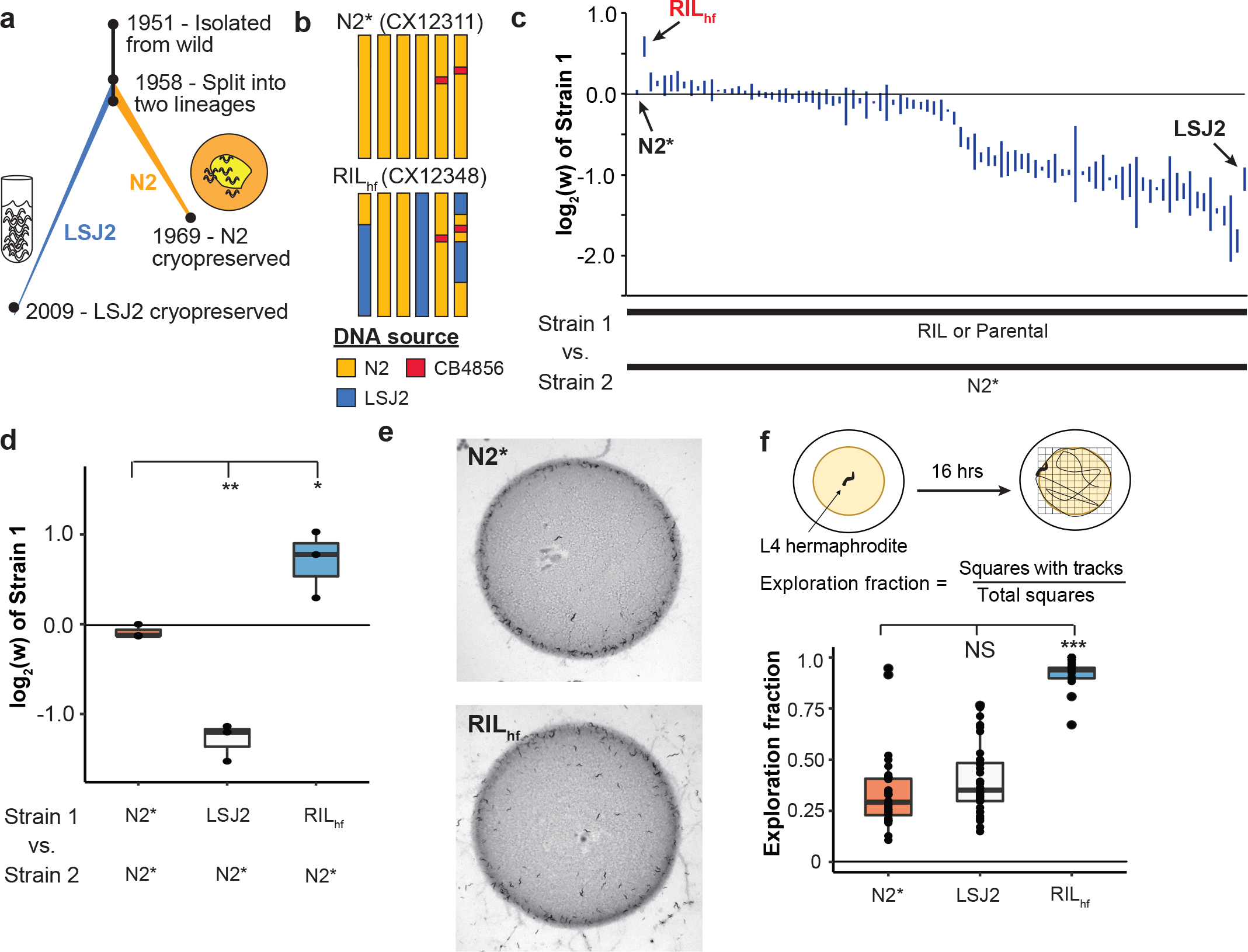
An outlier recombinant inbred line (RIL) showed higher evolutionary fitness and foraging behavior than either parental strain. **a.** Overview of the life history of two laboratory strains of *C. elegans* since their isolation from the wild in 1951 and subsequent split into two separate lineages around 1958. The standard reference N2 strain was cultured on agar plates seeded with *E. coli* bacteria until methods of cryopreservation were developed. LSJ2 was cultured in liquid, axenic media until 2009 when a sample of the population was cryopreserved. Resequencing of these strains identified ~300 genetic differences that fixed in one of the two lineages. **b**. Schematic of two strains used throughout this report. N2* (or CX12311) is a near-isogenic line (NIL) containing ancestral alleles of two genes, *glb-5* (chromosome V) and *npr-1* (chromosome X) backcrossed from the CB4856 wild strain. Beneficial alleles in these two genes fixed in the N2 lineage; use of the N2* strain allows us to exclude the effects of these alleles from our studies. RIL_hf_ (or CX12348) is an RIL strain generated between N2* and LSJ2, highlighted for its unexpectedly high levels of evolutionary fitness as determined in later experiments. RIL_hf_ contains LSJ2 sequence on chromosome IV and parts of the chromosome I and the X chromosome. **c**. Relative fitness levels were measured for a panel of 89 RIL strains generated between N2* and LSJ2 by competing each RIL against N2* for five generations. RILs were ordered by their average fitness value (three replicates were performed for each). Parental strains were also assayed (N2* and LSJ2). RIL_hf_ (red) is highlighted for its unusually high fitness, comparable to the fitness advantages of the N2 alleles of *glb-5* and *npr-1.* **d**. The fitness advantage of RIL_hf_ to the N2* strain was verified in an independent experiment. **e**. In normal growth conditions, agar plates seeded with *E. coli* bacteria, RIL_hf_ animals are more likely to be found in the center or outside of the bacterial lawn than at the borders. **f**. Foraging behavior differences were quantified by placing a single animal on a plate seeded with a circular lawn. After 16 hours, the amount of the plate that was explored was quantified by counting the number of grid squares with animal tracks within it (each point is a single animal). The RIL_hf_ explored more of the plate than either parental strain.

To determine if any other fixed mutations in the N2 lineage could affect fitness in standard laboratory conditions, we used a previously described panel of 89 recombinant inbred lines (RILs) between the CX12311 and LSJ2 strains (McGrath, Xu *et al.* 2011). CX12311 is a near isogenic line that carries ancestral *npr-1* and *glb-5* alleles from the CB4856 Hawaiian wild strain introgressed into an N2 background (**Figure 1b** - henceforth referred to as N2*) (McGrath, Xu *et al.* 2011). Using N2* as a parental strain eliminates the fitness effect of the derived alleles of N2 *npr-1* and *glb-5* (Zhao, Long *et al.* 2018). For each of the RIL strains, along with their parental lines, we performed pairwise competition experiments against a barcoded N2* strain to measure the relative RIL fitness compared to the common strain (Figure 1c **and Figure S1**). Interestingly, we found that one of the 89 RILs, CX12348 (henceforth called RIL_hf_ – hf for <u>h</u>igh <u>f</u>itness), had significantly higher fitness than either of the LSJ2 or N2* parental strains, which we validated in an independent experiment (**Figure 1d**). RIL_hf_ contained a mixture of DNA from both the N2* and LSJ2 parental strains, with N2* DNA on the left arm of chromosome I, the entire chromosomes II, III, and V, and portions of the X chromosome (**Figure 1b**). We decided to focus on this unusual strain_f_ to determine the genetic basis of its higher fitness, reasoning that a *de novo* mutation that occurred during construction of the RILs could be responsible for the transgressive phenotype of the RIL_hf_ strain.

Wild strains of *C. elegans*, as well as the N2* and LSJ2 parental strains, feed in groups on the borders of bacterial lawns, a foraging strategy known as social behavior (de Bono and Bargmann 1998). While growing the RIL_hf_ strain on standard plates, we noticed that animals had a stronger propensity to explore the centers and regions outside of the bacterial lawns, causing an increased number of worms and tracks in the center and outside of the lawn (**Figure 1e**). To quantify this behavioral difference, we modified a previously described exploration assay (Flavell, Pokala *et al.* 2013) to measure long-term (16h) exploration behavior in the presence of circular lawns (instead of uniform lawns). The exploration assay is an indirect measure of the relative time *C. elegans* spends in roaming and dwelling states in the presence of O_2_ gradients (and other chemical gradients) created by the circular bacterial lawns that are known to modify foraging behaviors (de Bono, Tobin *et al.* 2002, Gray, Karow *et al.* 2004). The RIL_hf_ strain explored a substantially larger fraction of the bacterial lawn than either of the parental strains (**Figure 1f**).

This change in foraging behavior is potentially an adaptive strategy for RIL_hf_ animals to increase their evolutionary fitness in the laboratory, or it might be a pleiotropic effect of the underlying genetic basis of this fitness gain. Since this foraging trait is easier to assay than relative fitness, we first focused on mapping this phenotype. We created two new small panels of 48 RILs between RIL_hf_ and either the N2* or LSJ2 parental strains and measured their foraging behavior (**Figure 2a, b**). An approximately equal number of RIL strains showed each parental phenotype, suggesting this trait was controlled by a single locus. We grouped these strains into low or high exploration groups for each RIL panel and performed pooled genomic sequencing on the four groups that were created (**Figure 2c**). A large allelic imbalance between the high and low exploration groups was observed in the center of chromosome III in the RIL_hf_ x LSJ2 panels (**Figure 2d**). This result is expected if a *de novo* mutation arose and fixed in the RIL_hf_ strain in the center of chromosome III (which contains the N2 haplotype for the entire chromosome). In this scenario, a similar imbalance would occur in the RIL_hf_ x N2* cross, however, because the two strains are largely identical on chromosome III, we could not observe it using the LSJ2/N2 SNVs.

**Figure 2.**
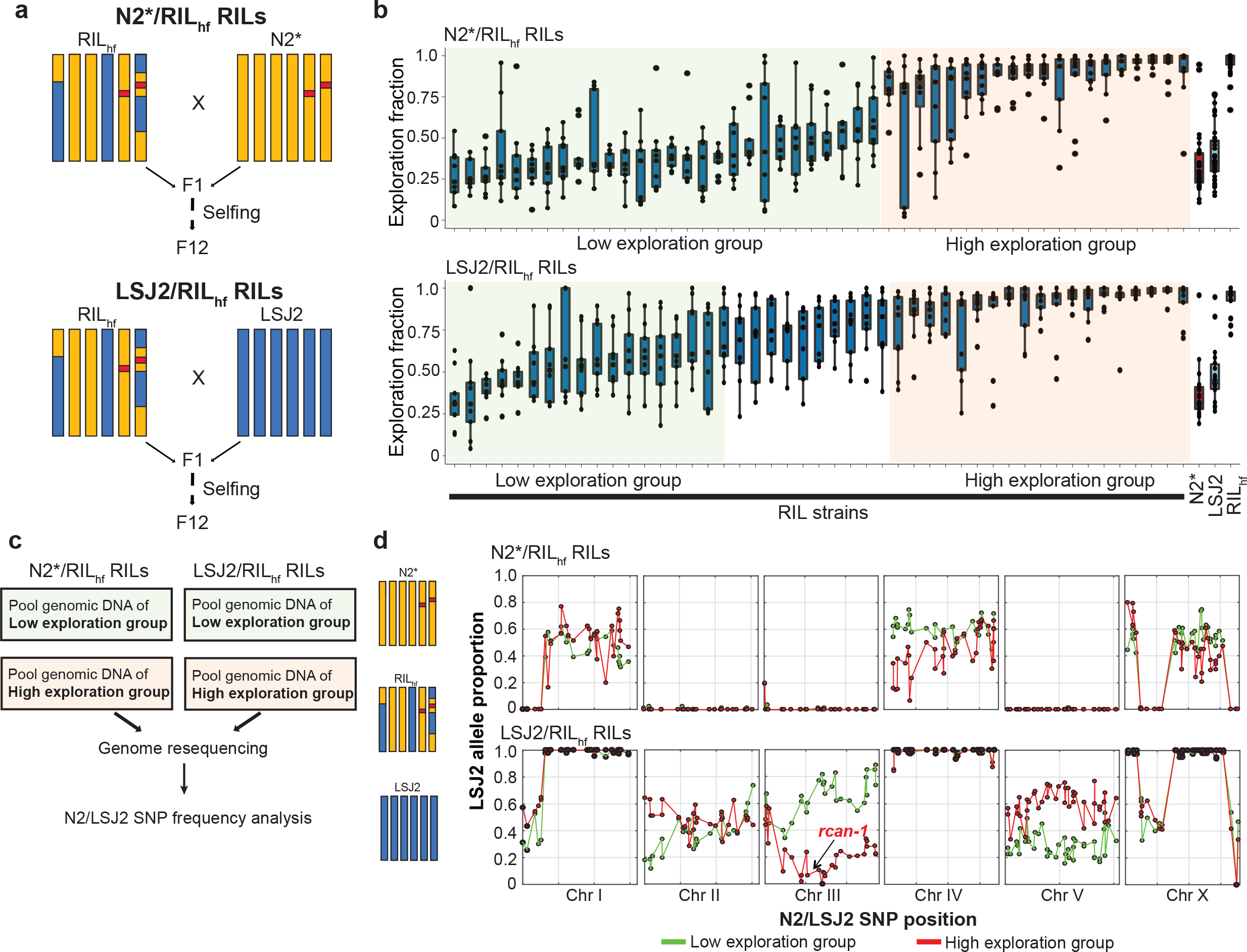
Foraging differences of the RIL_hf_ strain maps to the center of chromosome III. A. To map the changes in RIL_hf_ foraging behavior and fitness, we generated two panels of RILs (n = 48) between RIL_hf_ and N2* or RIL_hf_ and LSJ2. **b**. Each of the new RILs was measured for foraging behavior (shown in **Figure 1f**). An approximately equal number of low (green) and high (orange) exploration RILs was found, suggesting a single locus was responsible for the foraging differences. **c**. Overview of bulk-segregant approach from pooling genomic DNA isolated from the RILs into four samples. **d**. The four DNA samples were resequenced and the allele frequency for LSJ2/N2 genetic differences was estimated for each population. A large allelic frequency difference was observed on chromosome III in the LSJ2/RIL_hf_ RIL population between the low (green) and high (red) exploration groups.

To determine if the RIL_hf_ strain contains any *de novo* mutations in this region, we sequenced the RIL_hf_, N2*, and LSJ2 strains using Illumina short read sequencing. Although we did not identify any *de novo* SNVs or small indels on chromosome III in the RIL_hf_ strain, we did identify a large increase in coverage (2x – 8x) in the *rcan-1* gene, which is located in the center of chromosome III (**Figure 3a**). This coverage increase was detected in the high exploration groups of both RIL panels, consistent with this genetic change causing exploration behavior (**Figure 3a**).

**Figure 3.**
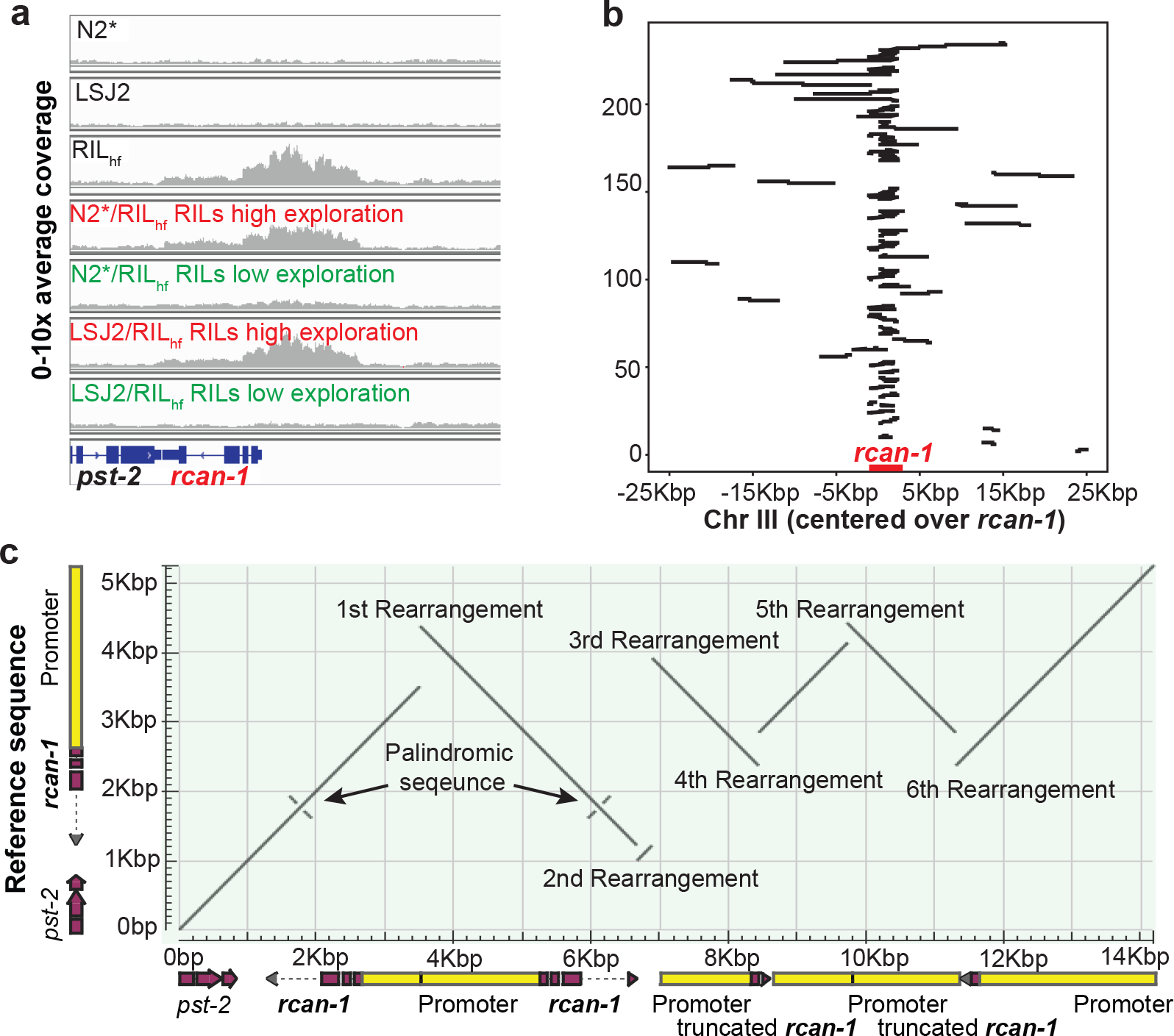
A de novo, complex rearrangement in *rcan-1*exists in the RIL_hf_ strain. **a.** The RIL_hf_ strain was re-sequenced to find *de novo* mutations on chromosome III that were not found in either the N2* or LSJ2 parental strains. A two- to eight-fold increase in coverage was observed across the *rcan-1* gene in the RIL_hf_ strain but not in either parental strain (each plot is normalized to 10x average coverage). This coverage increase was also found in the high exploration group of both RIL panels in **Figure 2**, consistent with this mutation causing the changes in foraging behavior. We were unable to resolve the exact nature of the genetic change from these data. **b**. The RIL_hf_ strain was sequenced using an Oxford Nanopore Minion device. Reads that mapped to *rcan-1* and containing a gap in alignment are shown. Alignment gaps can be caused by either poor sequence quality of the read, or by genomic rearrangements in the RIL_hf_ strain. The x-axis shows the position of the read relative to *rcan-1*, and the y-axis shows each contiguous mapping (*i.e.* each read spans two or more y-units). A single read (top) spanned the entire rearrangement (from −10 kb to 15 kb). **c**. The exact nature of the rearrangement was resolved using a combination of Nanopore and Illumina reads, plotted as a dot plot. The y-axis shows the reference sequence of *rcan-1*, and the x-axis shows the rearrangement in the RIL_hf_ strain. A total of six new junctions was observed, causing changes to the *rcan-1* locus shown under the x-axis.

While a simple gene duplication event would cause an increase in coverage, we observed a nonuniform change in coverage across the affected region. We also identified a large number of chimeric reads (reads which partially align to two unique locations) that mapped to five unique locations within the *rcan-1* locus (**Figure S2**). Together, these results suggest that the genetic change consisted of multiple inversion and/or duplication events. To resolve the precise mutation, we first attempted to amplify the entire affected region using PCR without success. As an alternative approach, we sequenced the RIL_hf_ strain using an Oxford Nanopore sequencing MinION, a long-read single molecule sequencing device with reported read lengths that could resolve the complex rearrangement (Tyson, O’Neil *et al.* 2018). By selecting reads that mapped to the *rcan-1* region, we identified a single 34,135 nucleotide long read that spanned the entire *rcan-1* region (**Figure 3b** and **S3**). By combining the Illumina short read, targeted PCR of portions of the rearrangement, and Nanopore long read data, we resolved the complex rearrangement into five unique tandem inversions interspaced within the *rcan-1* locus (**Figure 3c**, **S4**, and **File S1**).

To determine if this rearrangement was responsible for the increases in exploration behavior and relative fitness of the RIL_hf_ strain, we created two near isogenic lines (NILs) by backcrossing the *rcan-1* rearrangement from the RIL_hf_ strain into the N2* background. Genomic DNA from these NILs was sequenced to confirm that LSJ2-derived DNA and RIL_hf_-specific mutations besides the rearrangement were removed from both NILs (**Table S1**). As expected, both of these NILs explored a higher fraction of the bacterial lawn (**Figure 4a**). Pairwise competition experiments between the NILs and the N2* strain also demonstrated that this rearrangement is associated with the increases in fitness (**Figure 4b**) as well as growth rate (**Figure 4c**).

**Figure 4.**
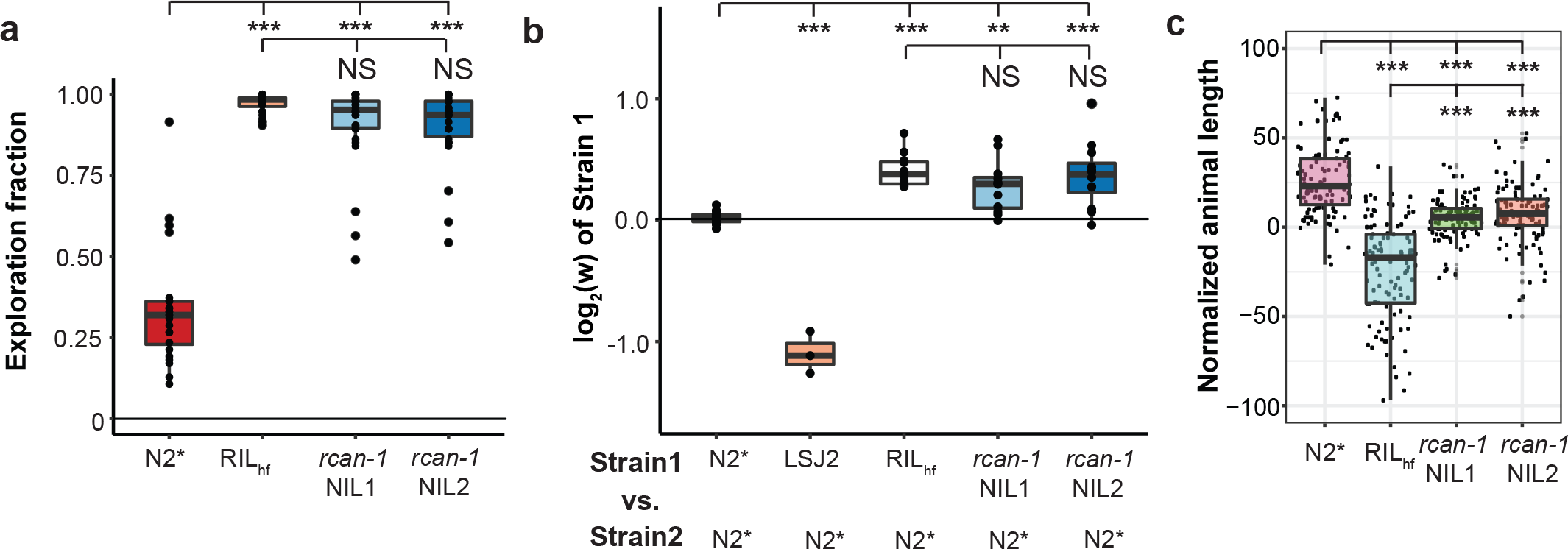
The *rcan-1*rearrangement is correlated with foraging differences and fitness gains of the RIL_hf_. **a.** We created two near isogenic lines (NILs) by backcrossing the RIL_hf_ strain to the N2* strain to remove all other LSJ2 and *de novo* variation from the RIL_hf_ strain. The two NIL strains showed a similar exploration fraction to the RIL_hf_ strain. **b**. Competition experiments between the NILs and N2*. Competition strains are shown on the x-axis, and the allele frequency of strain 1 is shown on the y-axis. The relative fitness differences of the NILs is comparable to the RIL_hf_ strain. **c**. A high-throughput assay was used to measure animal growth rates for N2*, RIL_hf_, and both backcrossed NILs. Each point is a biological replicate, with the y-axis indicating the median length of a population of animals. The animal sizes of both NILs are between the animal sizes of the parental strains.

The *rcan-1* rearrangement is predicted to cause a number of changes to the *rcan-1* gene. First, it creates two full-length versions of the *rcan-1* coding region (**Figure 3c**). However, the promoter region for each is modified by an inversion event in between the two coding regions. The first and second copies of *rcan-1* contain 857 and 1,725 bp of endogenous promoter before the inversion event occurs. Given that the nearest gene is over 5 kb upstream of the *rcan-1* gene in wild-type animals, enhancers and other regulatory regions of the promoter might be missing in the rearranged region, which might cause decreased, increased, or ectopic expression of *rcan-1.* Second, the second copy of the *rcan-1* gene also contains a small inversion in the 3’ UTR region. This inversion could modify binding sites for small RNAs or other RNA-binding proteins that regulate the stability or translation of the mRNA product. Additionally, the 3’ end of the small inversion is fused to an upstream promoter region, consequently, the native transcriptional terminator is missing from the second full-length copy of *rcan-1.* Finally, two truncated copies of the *rcan-1* gene are also created, containing the first two exons of the gene. It is possible that these products are translated into small fragments of RCAN-1 protein fused to novel protein sequence that could modify cellular function. It is difficult to predict *a priori* which of these changes alone or in combination could cause the changes to foraging behavior and/or evolutionary fitness.

To gain insights into the transcriptional changes caused by the rearrangement, we used RNA-seq to compare the genome-wide expression differences between the two NIL strains and the N2* strain. Interestingly, the gene with the largest change in expression was *rcan-1*, indicating that the rearrangement <u>decreased</u> transcription of the *rcan-1* gene by about 75% (**Figure 5a**, **S5**, and **Table S2**). Because the transcriptional profiling only reports whole-body changes in total *rcan-1* expression, we created promoter fusions of fluorescent proteins to determine how the two modified promoters for the full-length *rcan-1* gene affected expression in the *rcan-1* rearrangement. We cloned the entire region between the two full-length versions of *rcan-1* in both directions to create a P_*rcan-1-R1*_∷mCherry construct (reporting expression of the first full-length version of *rcan-1* in the complex rearrangement) and a P_*rcan-1-R2*_∷mCherry construct (reporting expression of the second full-length version of *rcan-1* in the complex rearrangement). As a control, we also cloned the first 5,085 bp of the wild-type promoter from N2 and fused it to both GFP and mCherry (P_*rcan-1-WT*_∷GFP or P_*rcan-1-WT*_∷mCherry). We then simultaneously co-injected P_*rcan-1-WT*_∷GFP with P_*rcan-1-WT*_∷mCherry, P_*rcan-1-R1*_∷mCherry, or P_*rcan-1-R2*_∷mCherry (**Figure 5b**). We used confocal microscopy to image whole-body expression from both green and red channels (**Figure S6**). As expected from a previous publication, we observed wild-type expression of *rcan-1* in a variety of tissues, including neurons, pharyngeal cells, and hypodermal cells. We first measured how the modified promoters affected whole-body expression by measuring the total amounts of GFP and mCherry signals from ~30 animals for each promoter construct (**Figure 5b**). Both of the promoters from the complex rearrangement drove less mCherry expression than the wild-type promoter to different extents, with the first promoter rearrangement more affected. The effect of the rearrangement was also tissue-specific, which we measured by quantifying the appropriate anatomical regions. For example, the head expression was significantly more affected then the body expression in both promoters (**Figure 5b**). The most prominent example is the approximately wild-type expression of two neurons in the retrovesicular ganglion that send a single process to the nerve cord (**Figure S7**). We tentatively identified these neurons as RIF or RIG. Combined with the whole body RNA-seq data, we suggest that *rcan-1* expression is largely reduced in the RIL_hf_ strain because of tissue-specific reductions driven by changes to the promoter regions of *rcan-1*.

**Figure 5.**
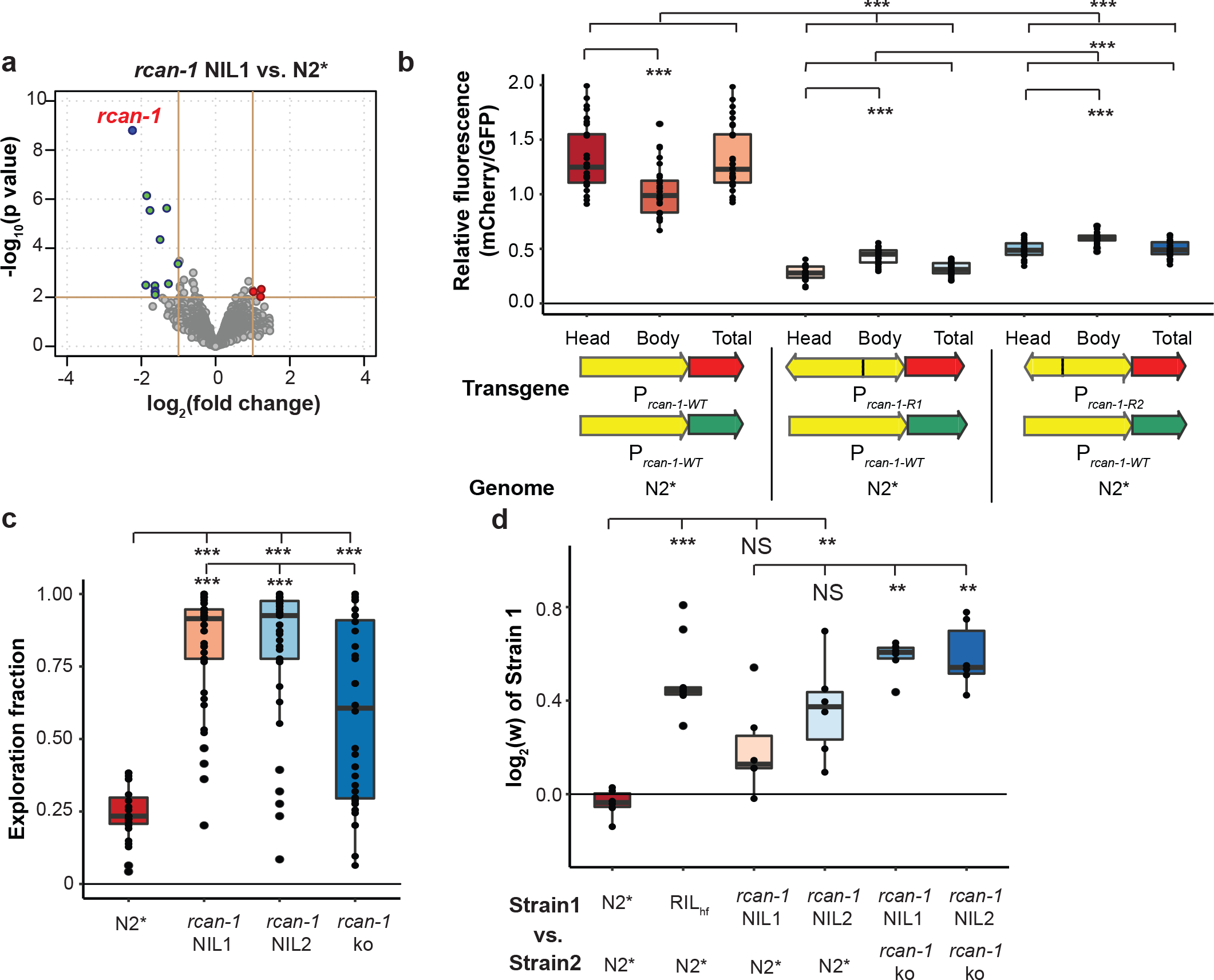
The *rcan-1* rearrangement allele causes reduced, tissue-specific expression and is not a loss-of-function allele. **a.** A volcano plot of *rcan-1* NIL expression vs. N2* animals is shown. Animals were synchronized, and RNA was isolated from L4 animals. The gene with the largest and most significant expression change was *rcan-1*, indicating the rearrangement causes reduced expression. **b**. Co-injection of wild-type *rcan-1* promotors driving GFP with wild-type *rcan-1* or rearranged *rcan-1* promoters created from RIL_hf_ driving mCherry were used to characterize how the rearrangement affected expression of the two full-length copies of the *rcan-1* coding region. Each dot represents the ratio of total GFP expression divided by total mCherry expression from a single animal. Fluorescence was also segmented into head or body expression and compared separately. Both rearranged promoters lowered expression. **c**. A large deletion of the *rcan-1* coding region was created using CRISPR-enabled genomic editing of the N2* strain. The *rcan-1* knockout modified foraging behavior but did not phenocopy the *rcan-1* NIL strains. **d**. Competition experiments demonstrated that the *rcan-1* deletion allele was less fit than the *rcan-1* rearrangement.

The above experiments suggest that the *rcan-1* rearrangement could be beneficial because of a reduction or loss of transcription in specific tissues while retaining wild-type or sufficient transcription in other tissues. However, an alternative hypothesis is simply that the rearrangement is beneficial because loss of *rcan-1* activity is beneficial in laboratory conditions and the remaining residual expression is unrelated to the fitness of the animals. To test the second hypothesis, we used CRISPR-enabled genomic editing to delete *rcan-1* in the N2* strain (**Figure S8**). This knockout strain showed an intermediate phenotype between the N2* and the *rcan-1* NIL strains in the modified foraging behavior (**Figure 5c**). When we competed this strain against the two *rcan-1* NILs, we found that the *rcan-1* rearrangement was substantially more fit than the *rcan-1* deletion (**Figure 5d**). We suggest that the complexity of the rearrangement is necessary for the fitness increases.

*rcan-1* is an ortholog of the human *RCAN1* gene (Li, Bell *et al.* 2015), which encodes a calcipressin family protein that inhibits the calcineurin A protein phosphatase (Fuentes, Genesca *et al.* 2000). In mammals, the exact genetic dosage of *RCAN1* is important; it has been proposed to be a key contributor to Down Syndrome phenotypes in patients with trisomy 21 (Fuentes, Genesca *et al.* 2000, Arron, Winslow *et al.* 2006). Additionally, chronic overexpression of *RCAN1* in mice promotes a number of mutant phenotypes related to Alzheimer’s disease (Martin, Corlett *et al.* 2012). In *C. elegans*, less is known about the function of *rcan-1.* It has been shown to be required for memory of temperature exposure through a *tax-6*/calcineurin-family and *crh-1*/CREB-dependent pathway (Li, Bell *et al.* 2015). Thermotaxis, however, is not predicted to be important for laboratory fitness, and it is likely that the *rcan-1* rearrangement regulates other unknown aspects of *C. elegans* biology on which selection can act. In mice, it was recently shown that *RCAN1* regulates weight gain on high fat diets, suggesting that *RCAN1* regulates energy expenditure (Rotter, Peiris *et al.* 2018). If this function is conserved in *C. elegans*, the excess energy extracted from abnormally high food levels of laboratory conditions might be diverted towards traits that improve evolutionary fitness. Unlike the standard N2 strain, which is potentially more fit in laboratory environments due to its ability to consume more food than the N2* strain, the *rcan-1* rearrangement showed no effect on food consumption (**Figure S9**).

Gene duplicates are thought to be a primary source of genetic material for the generation of evolutionary novelty, however, it is unclear how duplicates can arise and then navigate an evolutionary trajectory from redundancy to a state where both copies are maintained by natural selection as paralogs (Innan and Kondrashov 2010). One of the major issues in understanding how gene duplicates spread is understanding when divergence begins and what types of mutations lead to divergence. In one scenario, it is assumed that some sort of genetic event creates two copies of a gene that fix because of genetic drift or natural selection on redundancy (Clark 1994, Lynch and Force 2000). Adaptive evolution studies of nutrient limitations have also shown that gene duplicates can spread through positive selection, provided they facilitate the metabolism or transport of the limiting nutrient (Brown, Todd et al. 1998, Dunham, Badrane et al. 2002, Gresham, Desai et al. 2008, Kao and Sherlock 2008). Once fixed, functional diversification of the two gene copies, either through changes in gene expression or protein function, must occur for selection to maintain both in a population. However, because most mutations are deleterious, it is predicted that the fate of most gene duplicates is the loss of one copy from the population (Lynch and Conery 2000). Extant paralogs could represent the exceedingly rare events where gene duplication has occurred, mutation in one or both copies creates subfunctionalization or neofunctionalization, and then both copies can be maintained by selection. An alternative hypothesis proposes that functional divergence on segregating alleles, generated through balancing selection, precedes duplication (Proulx and Phillips 2006).

Our results demonstrate that duplication and functional diversification can happen simultaneously. The rearrangement of *rcan-1* occurred in a 12-generation pedigree used to construct 96 RIL lines. We propose that this rearrangement occurred as a single event, potentially as a result of an error in Okazaki fragment processing during DNA replication. The rearrangement of *rcan-1* causes large-scale changes to promoter regions, 3’UTR regions, and the creation of truncated versions of the *rcan-1* gene. For short evolutionary timescales, this type of genetic variant could potentially access changes to cellular function that would be difficult for a single single-nucleotide, insertion-deletion, or copy-number variant to cause. For example, besides changing the exact levels and tissues *rcan-1* gene is expressed, it is also predicted to create an mRNA with a modified 3’ UTR, potentially decoupling translational regulation or modifying mRNA stability in a subset of tissues. From a macroevolutionary perspective, the generation of two full-length versions of a gene, each with different promoters and regulatory regions, provides a flexible substrate for potentially generating two paralogs with different functions. Further, the beneficial nature of this mutation will result in its spread through the population, creating a large number of copies for mutation to generate further functional diversity. This study suggests that functional diversity between two gene duplicates can occur in a single event, short-circuiting much of the evolutionary process necessary for the spread and retention of gene duplicates. Genetic changes like this complex rearrangement are still difficult to identify using standard resequencing technologies and bioinformatic algorithms. As long-read sequencing technology improves, it will be interesting to determine the extent that similar alleles to the *rcan-1* rearrangement segregate within populations and the extent of their contribution to the unique genomes, forms, and behaviors of extant species.

## MATERIALS AND METHODS

### Growth Conditions

Animals were cultivated on standard nematode growth medium (NGM) plates containing 2% agar seeded with 200 μL of an overnight culture of the *E. coli* strain OP50 (Brenner 1974). Ambient temperature was controlled using an incubator set at 20°C. Strains were grown for at least three generations without starvation before any experiments were conducted.

### Strains

The following strains were used in this study. For each figure, a list of strains used is included in **Table S3**.

### Near isogenic lines (NILs)

CX12311 (N2*) - *kyIR1(V, CB4856>N2), qgIR1(X, CB4856>N2)*

PTM413 (*rcan-1* NIL 1) *kahIR16(III, CX12348>N2), kyIR1(V, CB4856>N2), qgIR1(X, CB4856>N2)*

PTM414 (*rcan-1* NIL 2), *kahIR17(III, CX12348>N2), kyIR1(V, CB4856>N2), qgIR1(X, CB4856>N2)*

### Recombinant inbred lines (RILs)

*CX12311 – LSJ2 RILs:* CX12312-19, CX12321-27, CX12346-52, CX12354-60, CX12362-66, CX12368-75, CX12381-88, CX12414-37, CX12495-99, CX12501-08, CX12510, CX12361

*CX12311 - CX12348 (RIL_hf_) RILs:* PTM378-397, PTM421-434, PTM494-503

*LSJ2 - CX12348 (RIL_hf_) RILs:* PTM435-478

### CRISPR-generated knockout and barcoded strains

PTM505: *dpy-10 (kah83)* II, *rcan-1(kah183) III, kyIR1(V, CB4856>N2), qgIR1(X, CB4856>N2)*

PTM288: *kyIR1(V, CB4856>N2), qgIR1(X, CB4856>N2) dpy-10(kah83)II*;

### Extrachromosomal array strains

PTM553 *kyIR1(V, CB4856>N2), qgIR1(X, CB4856>N2), kahEx169[P_rcan-1-WT_∷GFP 25ng/μL; P_rcan-1-WT_∷mCherry 25ng/μL] Isolate 1.*

PTM554 *kyIR1(V, CB4856>N2), qgIR1(X, CB4856>N2), kahEx170[P_rcan-1-WT_∷GFP 25ng/μL; P_rcan-1-WT_∷mCherry 25ng/μL] Isolate 2.*

PTM555 *kyIR1(V, CB4856>N2), qgIR1(X, CB4856>N2), kahEx171[P_rcan-1-WT_∷GFP 25ng/μL; P_rcan-1-WT_∷mCherry 25ng/μL] Isolate 3.*

PTM556 *kyIR1(V, CB4856>N2), qgIR1(X, CB4856>N2), kahEx172[P_rcan-1-WT_∷GFP 25ng/μL; P_rcan-1-R2_∷mCherry 25ng/μL] Isolate 1.*

PTM557 *kyIR1(V, CB4856>N2), qgIR1(X, CB4856>N2), kahEx173[P_rcan-1-WT_∷GFP 25ng/μL; P_rcan-1-R2_∷mCherry 25ng/μL] Isolate 2.*

PTM558 *kyIR1(V, CB4856>N2), qgIR1(X, CB4856>N2), kahEx174[P_rcan-1-WT_∷GFP 25ng/μL; P_rcan-1-R2_∷mCherry 25ng/μL] Isolate 3.*

PTM559 *kyIR1(V, CB4856>N2), qgIR1(X, CB4856>N2), kahEx175[P_rcan-1-WT_∷GFP 25ng/μL; P_rcan-1-R1_∷mCherry 25ng/μL] Isolate 1.*

PTM560 *kyIR1(V, CB4856>N2), qgIR1(X, CB4856>N2), kahEx176[P_rcan-1-WT_∷GFP 25ng/μL; P_rcan-1-R1_∷mCherry 25ng/μL] Isolate 2.*

PTM561 *kyIR1(V, CB4856>N2), qgIR1(X, CB4856>N2), kahEx177[P_rcan-1-WT_∷GFP 25ng/μL; P_rcan-1-R1_∷mCherry 25ng/μL] Isolate 3.*

### Strain construction

To create the CX12311-CX12348 and LSJ2-CX12348 RILs, CX12311 males or LSJ2 males were crossed to CX12348 hermaphrodites. 96 F2 progeny (48 from CX12311 x CX12348; 48 from LSJ2 x CX12348) were cloned to individual plates and allowed to self-fertilize for 10 generations to create the inbred lines. One RIL line was lost from the LSJ2xCX12348 cross, creating 47 RILs.

To create the *rcan-1* NILs (PTM413 and PTM414), CX12348 animals were backcrossed to CX12311 for 10 generations. Two completely independent sets of crosses were used to create two independent lines. Primers used to identify male animals containing the rearrangement were: 5’ - gagacaatactctgatattagacgcacca −3’ and 5’ – gctgacaccagcaatcattgttca −3’.

To create the *rcan-1* deletion strain (PTM505), two sgRNAs targeting the 5’ region of *rcan-1* and two sgRNAs targeting the 3’ end of *rcan-1* were created: sgRNA1: 5’-tcaacgaaatccgttgccaa-TGG-3’; sgRNA2: 5’-agtgctgatcaatgatccat-TGG-3’; sgRNA3: 5’-cgtggcatttcaattgctga-TGG-3’; sgRNA4: 5’-tcacatggagatgaagggcg-TGG-3’. CoCRISPR (Arribere, Bell *et al.* 2014) was used to simultaneously edit the *dpy-10* and *rcan-1* genes using the following injection mix: 50ng/μL P*eft-3∷Cas9*, 10ng/μL *dpy-10* sgRNA, 25ng/μL of each of the four *rcan-1* sgRNAs, and 500nM *dpy-10(cn64)* repair oligonucleotide. This mix was injected into CX12311 animals and Dpy or Rol animals were singled and genotyped using PCR. An animal with the deleted sequence 5’-caatggatcattgatca…‥cacgcccttcatctccat-3’ was identified.

To create the GFP/mCherry extrachromosomal lines (PTM553-PTM561), four constructs were created. *P*_*rcan-1-WT*_∷GFP was created by amplifying the *rcan-1* promoter from CX12311 genomic DNA using primers 5’-ctg<u>GGCCGGCC</u>tcggttcaaatacctcatgggaca-3’ and 5’-tt<u>GGCGCGCC</u>tttttgttgttaacttatagaaaaaatttcagcaacca-3’ and cloning it into the pSM-GFP backbone with restriction enzyme sites 5’-*Fse*I and 3’-*Asc*I. To create *P*_*rcan-1-WT*_∷mCherry, the *rcan-1* promoter was amplified from CX12311 genomic DNA using primers 5’-tcggttcaaatacctcatgggaca-3’ and 5’-tttttgttgttaacttatagaaaaaatttcagcaacca-3’ and a pSM-mCherry backbone was amplified using primers 5’-attttttctataagttaacaacaaaaaAcaagtttgtacaaaaaagcaggct-3 and 5’-ccatgaggtatttgaaccgaatagcttggcgtaatcatggtcat-3’. The two fragments were assembled using HI-FI assembly (NEB E5520S). To construct the *P*_*rcan-1-R1*_∷mCherry and *P*_*rcan-1-R2*_∷mCherry plasmids, 5’-tttttgttgttaacttatagaaaaaatttcagca-3’ and 5’-gaaacgaaacaaggtgggtcc-3’ or 5’-tttttgttgttaacttatagaaaaaatttcagca-3’ and 5’-agcggacccaccttgtttc-3’ were used to amplify the rearranged promoters from CX12348 genomic DNA. These PCR products were cloned into a pSM-mCherry backbone using HI-FI assembly. Concentrations of each plasmid are indicated for each strain in the strain description.

### Competition experiment

Competition experiments were performed as described previously (Zhao, Long *et al.* 2018). In brief, 9 cm NGM plates were seeded with 300 μL of an overnight *E. coli* OP50 culture and incubated at room temperature for three days. Ten L4 larvae from each strain were picked onto the plates and cultured for five days. Animals were transferred to identically prepared NGM plates and subsequently transferred every four days for five to seven generations. For each transfer, animals were washed off the plates using M9 buffer and collected into 1.5 mL centrifuge tubes. The animals were then mixed by inversion and allowed to stand for approximately one minute to settle adult animals. 50 μL of the supernatant containing approximately 1000-2000 L1-L2 animals were seeded onto fresh plates. The remaining animals were concentrated and used for genomic DNA isolation. Genomic DNA was collected every odd generation using a Zymo DNA isolation kit (D4071). To quantify the relative proportion of each strain, a Taqman assay was performed by digital PCR based approach with custom TaqMan probes (Applied Biosciences). Genomic DNA was digested with *Sac*I or *Eco*RI for 30 min at 37 ºC. The digested products was purified using a Zymo DNA cleanup kit (D4064) and diluted to approximately 1-2 ng/μL. Seven TaqMan probes were designed using ABI custom software that targeted the SNP WBVar00051876, SNP WBVar00601322, SNP WBVar00167214, SNP WBVar00601493, SNP WBVar00601538, *dpy-10 (kah82)*, or *tbc-10(kah185).* The Taqman digital PCR assays were performed using a Biorad QX200 digital PCR machine with standard probe absolute quantification protocol. The relative fitness was calculated using a haploid selection model (Zhao, Long *et al.* 2018). The relative fitness value and Taqman assay information for each competition experiment are included in **Table S3**.

### Exploration behavioral assay

The exploration assays from Flavell *et al.* were modified to study foraging in the presence of circular lawns (Flavell, Pokala *et al.* 2013). 35 mm Petri dishes were seeded with 150 μL OP50 *E. coli* Bacteria for 24 h before the start of the assay. Individual L4 hermaphrodites were placed in the center of the plate and cultivated in 20ºC for 16 hours. The plates were placed on a grid that has 100 squares that cover the whole bacteria lawn. To calculate the exploration fraction, the number of full or partial squares that contained animal’s tracks out of bacteria lawn border was quantified. The number of full or partial squares that contain the bacteria lawn was also counted (about 94-96 grids). The exploration fraction was calculated (equation 1).

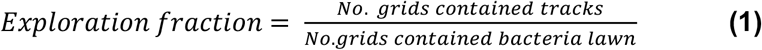

### Bulk-segregant analysis of exploration behavior

The exploration behavioral assays were performed on 48 CX12311-CX12348 RILs and 47 LSJ2-CX12348 RILs. In the CX12311/CX12348 RILs, 28 RILs with median exploration fraction less than 0.575 were assigned to the low exploration group and the 20 RILs with median exploration fraction greater than or equal to 0.575 were assigned to the high exploration group. In the LSJ2/CX12348 RILs group, the 17 RILs with median exploration fraction less than 0.620 were assigned to the low exploration group, the 20 RILs with median exploration behavior greater than or equal to 0.870 were assigned to high exploration group, and the rest of the RILs were excluded from further analysis. Genomic DNA from each RIL (100 ng) was isolated and pooled into the four described groups for whole-genome resequencing.

### Whole-genome resequencing

Genomic DNA was isolated using Qiagen Gentra Puregene Kit (158667) following the supplementary protocol for nematodes. The genomic DNA was further purified using Zymo Quick-DNA kit (D4068). DNA libraries were prepared using an Illumina Nextera DNA kit (FC-121-1030) with indexes (FC-121-1011). The prepared libraries were sequenced at 35 bp or 150 bp paired-read using an Illumina NextSeq 500. The reads were aligned to reference genome using BWA-aligner v0.7.17 (Li and Durbin 2009). BAM files were deduplicated and processed using SAMtools v1.9 (Li, Handsaker *et al.* 2009) and Picard. SNVs were called by Freebayes and annotated by SnpEff (Cingolani, Platts *et al.* 2012, Garrison and Marth 2012). Custom Python scripts using the pysam library (https://github.com/pysam-developers/pysam) were used to identify regions of the genome with a large number of clipped and chimeric reads. Reads depths were visualized using IGV (Robinson, Thorvaldsdóttir *et al.* 2011). The sequencing reads were uploaded to the SRA under BioProject PRJNA526525.

### Oxford Nanopore long-read sequencing

Genomic DNA of CX12348 was isolated from animals grown on 8 9 cm NGM plates using Qiagen Gentra Puregene Kit (158667) following the supplementary protocol for nematodes. The genomic DNA was concentrated and purified using Zymo Quick-DNA kit (D4068). Size-selection to collect DNA fragments from 10 kbp – 50 kbp was carried out using a Blue-pippin. The sequencing library was prepared using 1D ligation kit (SQK-LSK108) following the standard protocol. DNA was repaired using the NEBNext FFPE Repair Mix (M6630). After DNA repair, end preparation was performed and the adapter was ligated. 600ng prepared library was loaded in the Nanopore R9 flow cell in MinION sequencer. The standard 48 hours sequencing protocol was performed and approximately 5 Gb of sequencing data was generated. To resolve the structure of *rcan-1* complex rearrangement, the FASTQ files were aligned to reference genome using BWA aligner. Reads that covered the *rcan-1* gene region and contained a gap in alignment were fetched using pysam (https://github.com/pysam-developers/pysam). These reads were then mapped to *rcan-1* using BLAST and visualized with matplotlib (https://matplotlib.org) to show the rearrangement events. The structure of the complex rearrangement was verified by mapping Illumina short reads to the proposed structure. The sequencing reads were uploaded to the SRA under BioProject PRJNA526525.

### RNA-seq and transcriptome analysis

CX12311, PTM413, and PTM414 were synchronized using alkaline-bleach to isolate embryos, which were washed with M9 buffer and placed on a tube roller overnight. Approximately 400 hatched L1 animals were placed on NGM agar plates for each strain and incubated at 20°C for 48 hours. The ~L4 stage animals were washed off for standard RNA isolation using Trizol. Four replicates for each strain were performed on different days. The RNA libraries were prepared using an NEB Next Ultra II Directional RNA Library Prep Kit (E7760S) following its standard protocol. The libraries were sequenced by Illumina NextSeq 500. The reads were aligned by HISAT2 using default parameters for pair-end sequencing. Transcript abundance was calculated using HTseq and then used as inputs for the SARTools (Varet, Brillet-Guéguen *et al.* 2016). edgeR v3.16.5 was used for normalization and differential analysis. The analysis result was shown in a volcano plot. CX12311 was treated as the wild type (Chen, Lun *et al.* 2014). The genes show significant differential expression in the volcano plot are under thresholds |log_2_(fold)| > 1 and FDR adjusted p-value < 0.01. Sequencing reads were uploaded to the SRA under BioProject PRJNA526525.

### Imaging

The detailed steps of microfluidic device fabrication were previously reported(Lee, Kim et al. 2014). For each experiment, about 100-150 animals were suspended in 1 mL of S Basal and delivered into using a syringe. Animals were immobilized using 1 mL of tetramisole hydrochloride (200 mM) (Sigma-Aldrich cas. 5086-74-8) in S Basal. Imaging were acquired on a spinning disk confocal microscope (PerkinElmer UltraVIEW VoX) with a Hamamatsu FLASH 4 sCMOS camera. Images of the animals were quantified using ImageJ. A region-of-interest (ROI) was drawn around the entire worm, and the mean intensity of the GFP and mCherry images were calculated across the ROI. Relative fluorescence intensity was calculated as (Mean Intensity of mCherry)/(Mean Intensity of GFP).

### Food consumption assay

The experimental method was described previously (Zhao, Long *et al.* 2018). In brief, the food consumption assay was performed using 24-well plates seeded with 20μL of freshly cultured OD600 of 4.0 (CFU ~ 3.2×10^9^/mL) *E. coli* OP50-GFP(pFPV25.1). The fluorescence signal of OP50-GFP was quantified by area scanning protocol using BioTek Synergy H4 multimode plate reader. The synchronized L4 animals were placed in the wells in the first five columns and the last column is used as control column. Each well was placed with 10 animals, and the plate was incubated in a 20°C incubator for 18 hours and the fluorescence signal was quantified again as the ending time point. The relative food consumption amount was calculated using the equations reported previously (Zhao, Long *et al.* 2018).

### High-throughput growth rate analysis

The high-throughput growth rate and brood size assays were performed as described previously (Evans, Brady *et al.* 2018). In short, approximately 25 bleach-synchronized embryos were aliquoted into each well of 96-well plates, and fed 5 mg/mL HB101 bacterial lysate on the following day (Garcia-Gonzalez, Ritter et al. 2017). After 48 hours of growing at 20°C, a large-particle flow cytometer (COPAS BIOSORT, Union Biometrica, Holliston, MA) was used to sort three L4 larvae into each well of a 96-well plate with 50 μL of K medium with HB101 lysate (10 mg/mL) and Kanamycin (50 μM). Animals were grown for 96 hours at 20°C and were then treated with sodium azide (50 mM in M9). Animal number (n) and animal length (time of flight, TOF) were measured by the BIOSORT. For each well, animal growth was measured as the median length of the population, and brood size was measured as the number of progeny per sorted animal. The experiments were replicated in two independent assays, and the linear model with the formula (phenotype ~ assay) was applied to normalize the differences among assays (Shimko and Andersen 2014).

### QTL mapping

The average of the log2(w) of each N2*/LSJ2 RIL was used as phenotype with 192 previously genotyped SNPs. R/qtl was used to perform a one-dimensional scan using marker regression on the 192 markers. The genome-wide error rate (p = 0.05) was determined by 1000 permutations test.

### Statistical analysis

The raw data are included in Table S3. To assess statistical significance, we performed one-way ANOVA tests followed by Tukey’s honest significant difference test to correct for multiple comparisons. NS: not significant; * : *p* < 0.05; **: *p* < 0.01; ***: *p* < 0.001.

## Acknowledgements

We thank the *Caenorhabditis* Genetics Center for strains, members of the Streelman and McGrath lab for discussions, and WormBase. This work was supported by NIH R01GM114170 (to P.T.M), a John N. Nicholson fellowship (to S.C.B), an NSF CAREER Award (to E.C.A.), and NIH NS096581, GM088333, AG056436 (to H.L.).

