## Supplemental Figures 1-8 for "A beneficial genomic rearrangement creates multiple versions of calcipressin in *C. elegans*"

### Figure S1

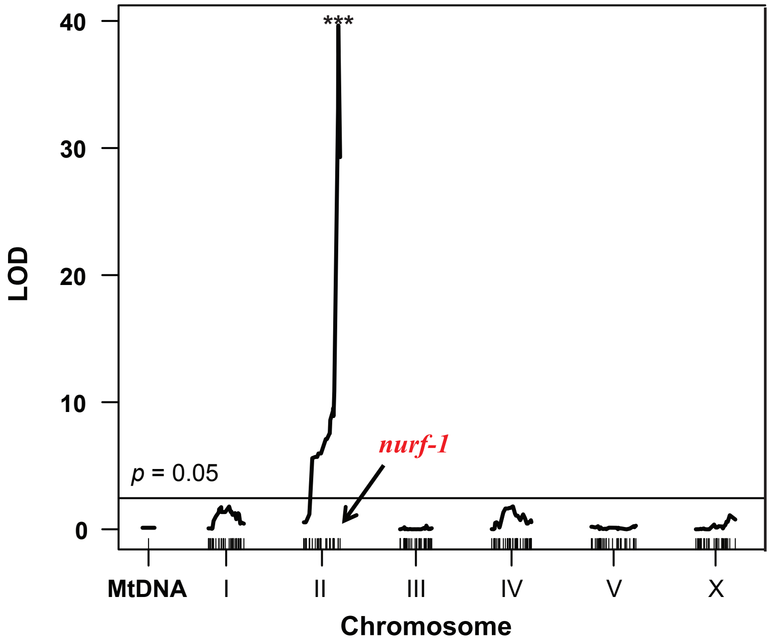


**Figure S1. QTL mapping of relative fitness.** We performed QTL mapping on the relative fitness differences between the RIL strains shown in Figure 1c. A single significant QTL on the right arm of Chomosome II, which overlaps the previously identified *nurf-1* gene, was identified. Threshold line is significance level at $p$ = 0.05 from a 1,000 permutation test.

### Figure S2

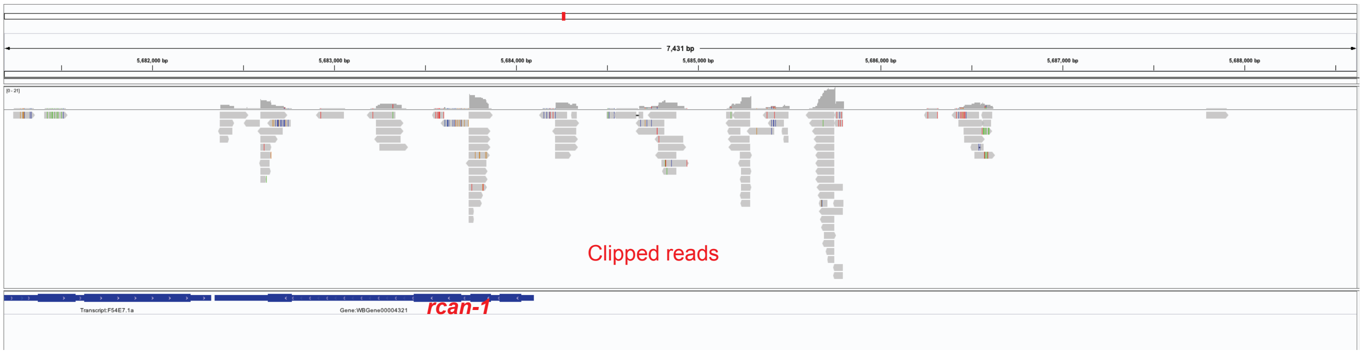


**Figure S2. Clipped reads mapped to the *rcan-1* locus.** An IGV plot of chimeric reads align to two genomic locations. Reads were from resequencing of the $RIL_{hf}$ (CX12348) strain.

### Figure S3

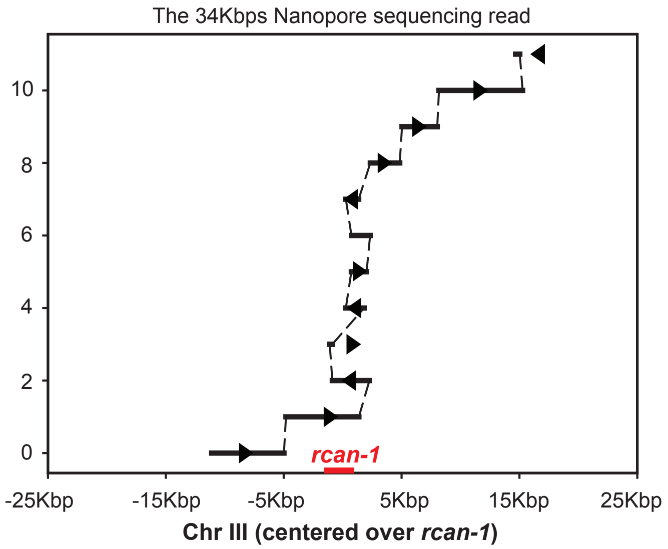


**Figure S3. A single Nanpore sequencing read resolves the *rcan-1* complex genomic rearrangement.** A 34 kbp Nanopore sequencing read was aligned to the to *rcan-1* reference sequence. The solid line shows alignment between the read and *rcan-1* as determined by blastn. The arrow indicates the alignment direction. Striped lines indicate gaps in the the alignment, which can be caused by either structural changes in the DNA (e.g. 1-2) or loss-of-quality in the Nanopore read (e.g. 0-1).

### Figure S4

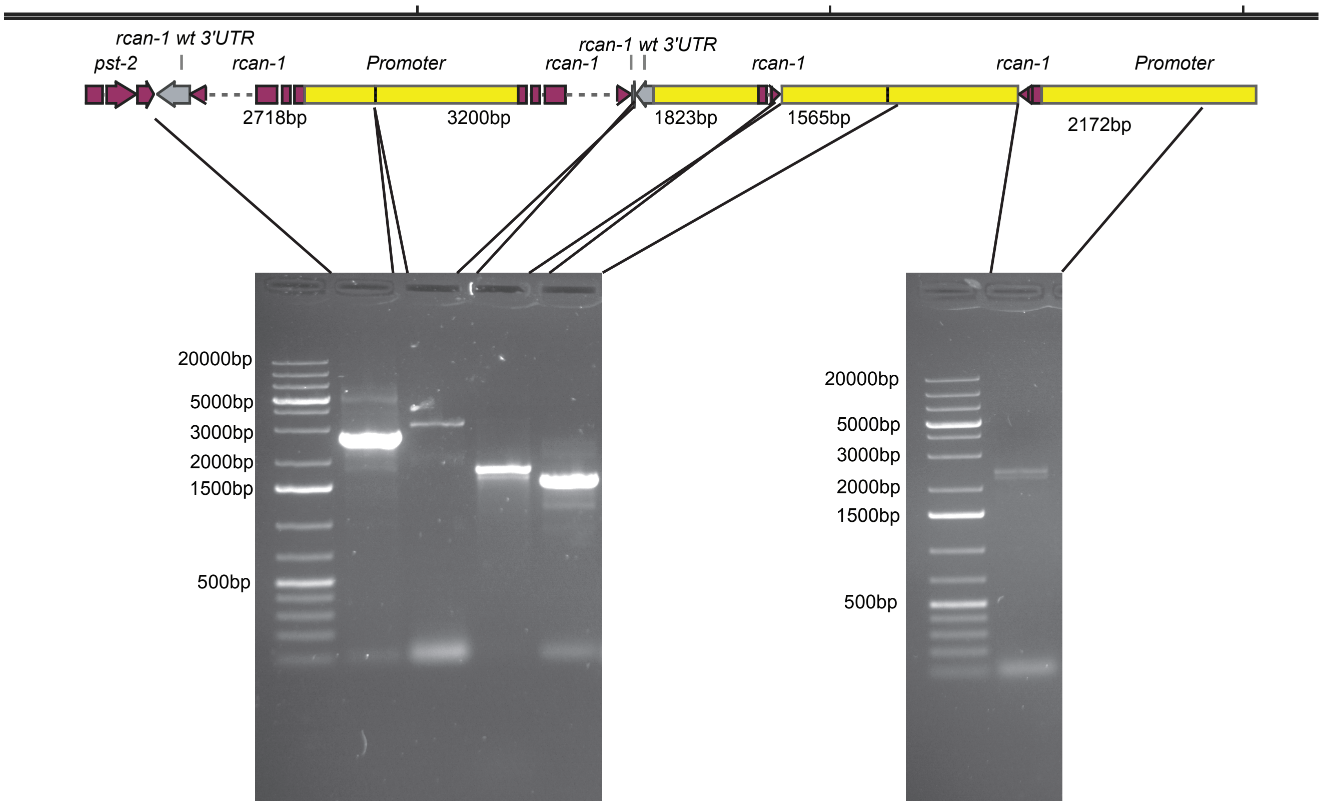


**Figure S4. PCR products that support the proposed *rcan-1* rearrangement.** Applicons for each lane were generated from primers that matched regions of *rcan-1* indicated by the black lines. These aplicons were sequenced to verify they amplify the expected locus.

#### Figure S5

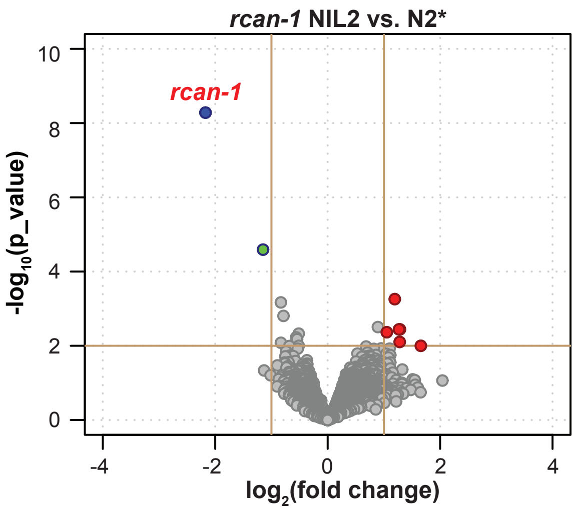


**Figure S5. Volcano plot of *rcan-1 NIL 2* gene expression vs. N2*.** Red dots indicate genes with increased expression in *rcan-1 NIL 2* vs. N2*. Green dots indicate genes with decreased expression in *rcan-1 NIL 2* vs. N2*.

### Figure S6

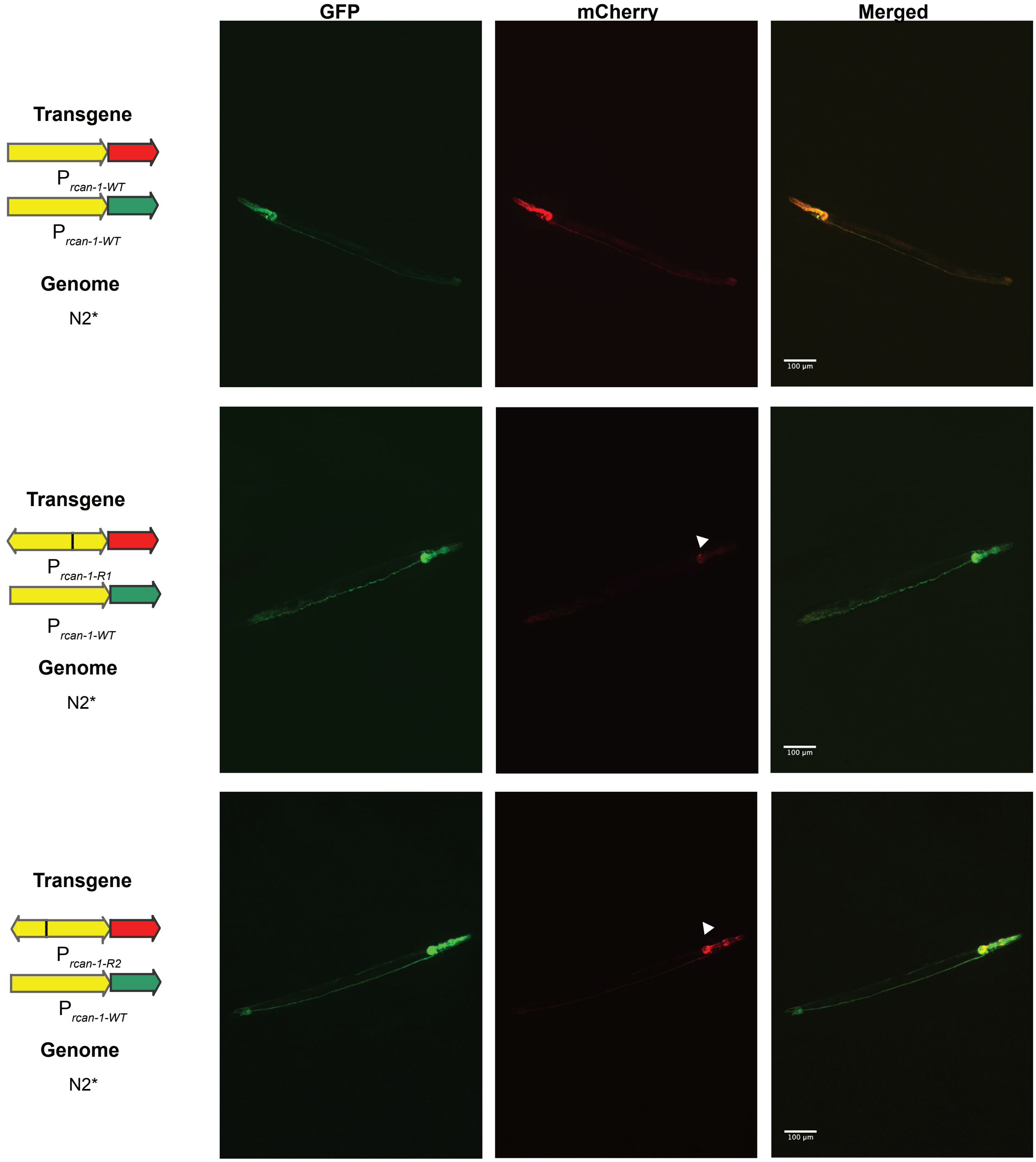


**Figure S6. Representative images of promoter constructs driving fluorescence reporters.** N2* animals were coinjected with wild-type *rcan-1* promotors driving GFP with either wild-type *rcan-1 (top)* or rearranged *rcan-1* promoters created from the $RIL_{hf}$ strain (middleand bottom) driving mCherry. The scale bar is 100um.

### Figure S7

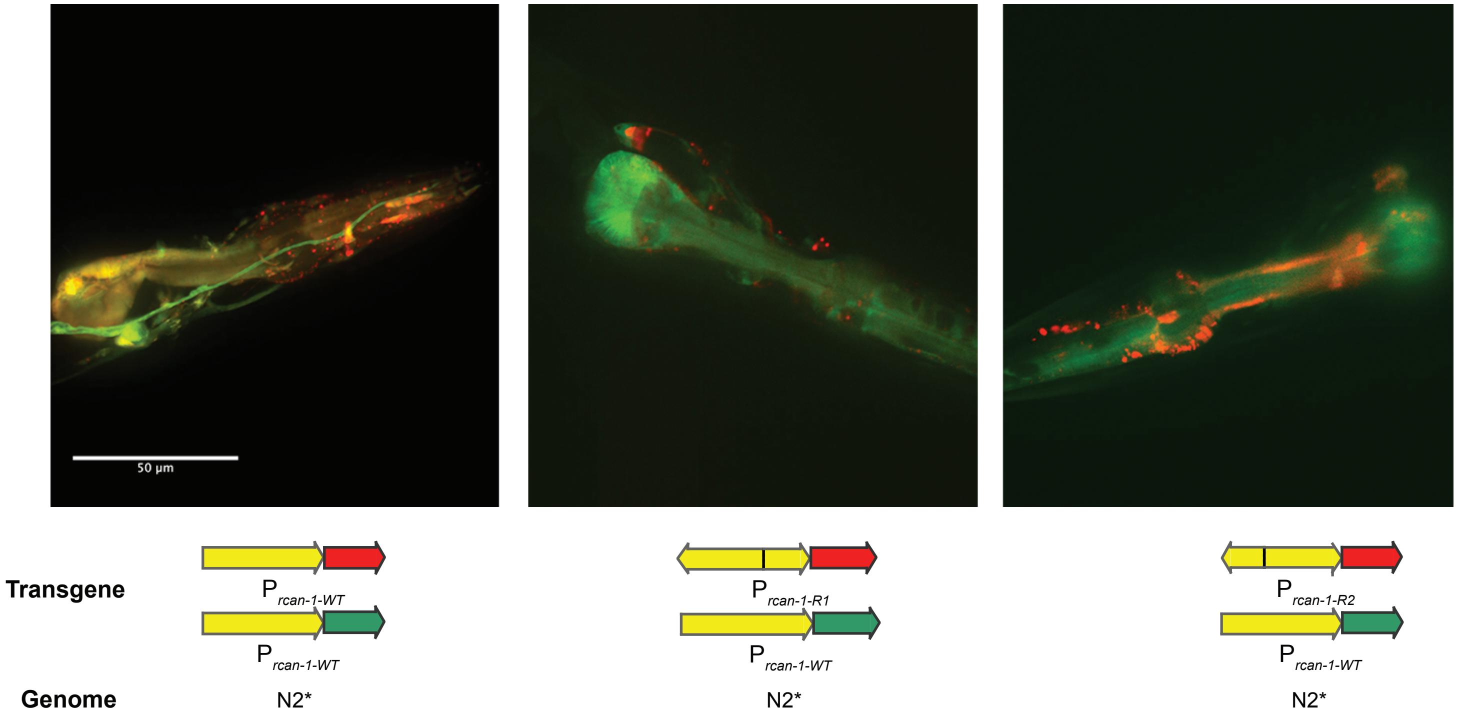


**Figure S7. Representative images showing tissue-specific differences in promoter expression.** N2* animals were coinjected with wild-type *rcan-1* promotors driving GFP with wild-type *rcan-1* or rearranged *rcan-1* promoters created from $RIL_{hf}$ driving mCherry. While expression in the pharynx is largely reduced in the rearranged promoters, expression of a pair of unknown interneurons was less affected. Scale bar is 50um.

### Figure S8

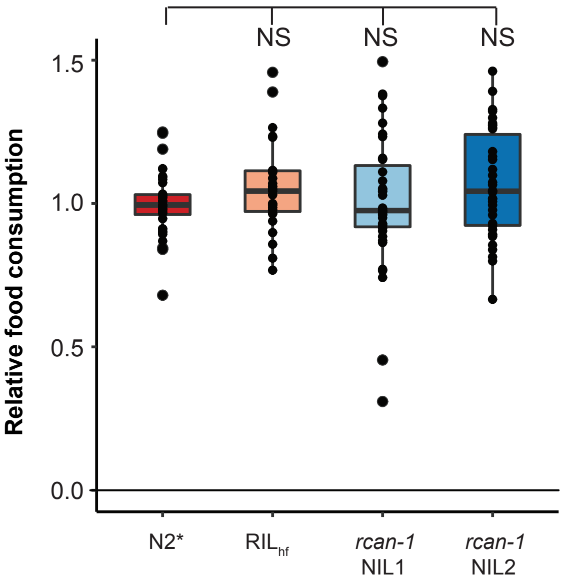


**Figure S8. Food consumption assay of RIL$_{hf}$ and *rcan-1 NILs*.** Relative food consumption of indicated strains. Each dot indicates one experimental replicate.
